# A comprehensive, FAIR file format for neuroanatomical structure modeling

**DOI:** 10.1101/2020.09.22.306670

**Authors:** A. E. Sullivan, S. J. Tappan, P. J. Angstman, A. Rodriguez, G. C. Thomas, D. M. Hoppes, M. A. Abdul-Karim, M. L. Heal, J. R. Glaser

**Affiliations:** MBF Bioscience, Williston, VT, USA

**Keywords:** Neuromorphology, Neuroimaging, Morphological modeling, Neuron reconstruction, FAIR data, Vasculature reconstruction

## Abstract

With advances in microscopy and computer science, the technique of digitally reconstructing, modeling, and quantifying microscopic anatomies has become central to many fields of biological research. MBF Bioscience has chosen to openly document their digital reconstruction file format, Neuromorphological File Specification (4.0), available at www.mbfbioscience.com/filespecification (Angstman et al. 2020). One of such technologies, the format created and maintained by MBF Bioscience is broadly utilized by the neuroscience community. The data format’s structure and capabilities have evolved since its inception, with modifications made to keep pace with advancements in microscopy and the scientific questions raised by worldwide experts in the field. More recent modifications to the neuromorphological data format ensure it abides by the Findable, Accessible, Interoperable, and Reusable (FAIR) data standards promoted by the International Neuroinformatics Coordinating Facility (INCF; Wilkinson et al. 2016). The incorporated metadata make it easy to identify and repurpose these data types for downstream application and investigation. This publication describes key elements of the file format and details their relevant structural advantages in an effort to encourage the reuse of these rich data files for alternative analysis or reproduction of derived conclusions.

## Background

Digitally reconstructing and modeling microscopic anatomies has become important in many fields of research, none more so than the field of neuroscience. One of the principle uses of this technique is 3D neuronal reconstruction, which allows researchers to create and analyze accurate and quantifiable neuronal models derived from microscopic specimens (Meijering 2010; Ascoli, Duncan, Donohue, and Halavi 2007). By geometrically representing the structures contained in the image data in a 3D volume, detailed morphometric analyses, simulations and electrotonic modeling of the neurons can be performed. Unlike raw microscopic image data, the reconstruction data specifies the individual neuronal components and denotes the X, Y, and Z position of every point of each of the modeled structures.

The field of digital morphological reconstruction has evolved and expanded for more than 50 years (Fig. 1). The origin of this technique was dates back to the seminal work of Edmund Glaser and Hendrik Van der Loos who first described it in the paper published in 1965, “A semi quantitative computer-microscope for the analysis of neuronal morphometry”. This paper describes a system for attaching X-Y-Z transducers to a microscope stage, tracing the branches of a Golgi-stained neuron, and outputting the result to a plotter (Fig. 1a; Glaser and Van der Loos 1965). This work was continued by the father and son team of Edmund Glaser and Jack Glaser, who founded MBF Bioscience (at that time known as MicroBrightField) in 1988. MBF Bioscience expanded on the original invention of Glaser and Van der Loos to develop Neurolucida (Fig. 1b; Glaser and Glaser 1990), which has become a widely used microscope system for neuron reconstructions in humans and other species (Halavi, Hamilton, Parekh, and Ascoli 2012; Parekh and Ascoli 2013; Blackman, Grabuschnig, Legenstein, and Sjöström 2014; Usher et al. 2018), with more than 6,500 citations (Rance 1990; Schiller, Schiller, Stuart, and Sakmann 1997; Wong, Wang, and Axel 2002; Henriksen, Colgin, Barnes, Witter, Moser, and Moser 2010; Földy, Malenka, and Südhof 2013; Lázaro, Hertel, Sherwood, Muturi, and Dechmann 2018; Ullah et al. 2020). Its popularity has overtaken other integrated microscopy systems such as the Eutectic Neuron Tracing System (NTS; Parekh and Ascoli 2013).

**Fig. 1.**
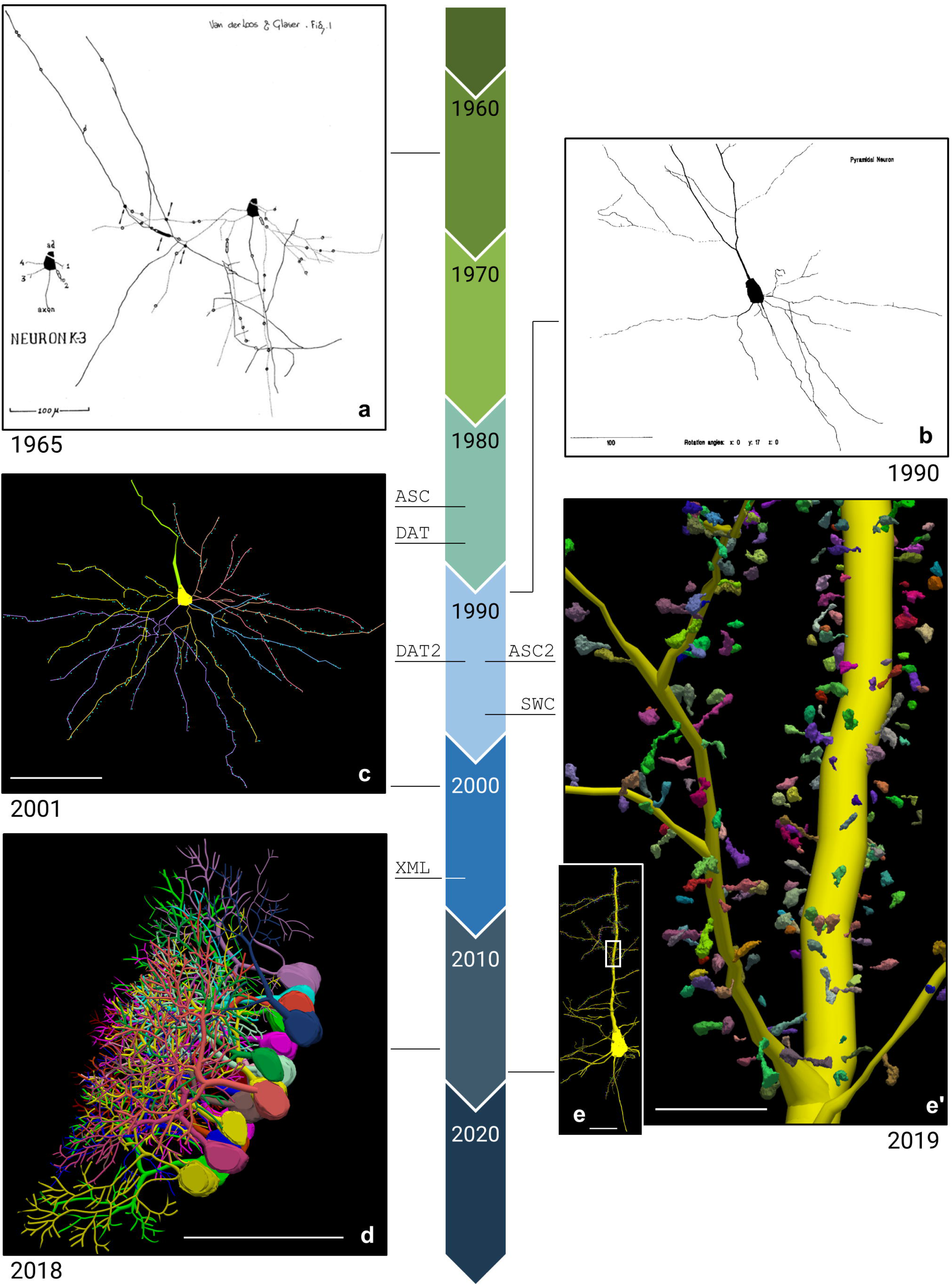
A timeline depicting the evolution of digital neuron reconstruction between 1960 and 2020. The lines connecting each image to the timeline indicate when in time the tracing was generated. The birth of each data file format is indicated with the line under each file name: ASC (1986), DAT (1988), DAT2 and ASC2 (1995), SWC (1998), XML (2007). The timeline’s colored arrows each represent one decade and are labeled with the first year of that decade. Each neuronal reconstruction includes the publication date below the image. **(a)** A neuronal reconstruction “produced by the first computer-assisted neuron tracing system, Neurolucida’s anacestor” (Glaser and Vanderloos 1965). The scale bar equals 100 micrometers. **(b)** “The hard copy, monochrome output of the neuron of Fig 3…**”** in the paper, “Neuron imaging with Neurolucida--a PC-based system for image combining microscopy.” (Glaser and Glaser 1990). The scale bar equals 100 micrometers. **(c)** A reconstruction of a human supragranular pyramidal cell from the Brodmann area (BA), superior frontopolar zone (BA10). Spines have been mapped using point markers (blue) along the cell’s dendrites (Jacobs 2001). The scale bar equals 100 micrometers. **(d)** Purkinje cells (PCs) reconstructed using Neurolucida 360 from the cerebellar vermis of male mice (Nedelescu, Abdelhack, Pritchard 2018). The scale bar equals 100 micrometers. **(e)** A Neurolucida 360 reconstruction of a *Drosophila* pyramidal neuron, showing the soma, apical/ basal dendrites, and axon segments (Gao et al. 2019). The white box demonstrates the approximate location of Fig. 1e’. The scale bar equals 100 micrometers. **(e’)** A zoomed in look at the 3D spines reconstructions of the cell tracing shown in Fig. 1e. The scale bar equals 25 micrometers.

From its initial release in 1988, Neurolucida made use of two file formats for storage and archival of tracing data: a portable ASCII text format, and a binary platform-dependent format. The ASCII text format, normally referred to as the Neurolucida ASC format, was conceived as a human-readable and portable format that could be easily imported into other software, regardless of underlying computing hardware or operating system. The binary format, referred to as the Neurolucida DAT format, was intended as a more computationally efficient alternative for use within the Neurolucida environment. Both of these formats have continued to evolve over the past 30 years to accommodate new research requirements and as of this writing are still fully supported by MBF products.

By the mid-1990’s, with the increase in memory and computing power of personal computers along with the connectivity afforded by the now burgeoning internet, full-cell neuronal tracings created with Neurolucida were being created and shared (Turner, Li, Pyapali, Ylinen, and Buzsaki 1995; Mpodozis, Cox, Shimizu, Bischof, Woodson, and Karten 1996; Jackson and Cauller 1997; Wu and Karten 1998; Prusky and Arjannikova 1999; Gabriele, Brunso-Bechtold, and Henkel 2000; Ghosh et al. 2011). In 1998, a seminal paper presented an online archive system for neuronal reconstructions (Cannon, Turner, Pyapali, and Wheal 1998). The archive, initially populated by a set of 87 neurons from the hippocampus reconstructed using Neurolucida, offered an editor and a format converter. The system’s native archival file format, SWC, was a simple and portable text-based format that could represent arbitrarily-complex neuronal geometries by providing a list of positions and radius in a hierarchical parent/child arrangement that form a collection of minimally connected cylindrical segments. Due to its simple, portable, and topologically constrained format, SWC became a preferred file format for archival and sharing of neuronal tracings for use in compartment modeling and computational neuroanatomy applications. One notable example of SWC’s success has been its adoption as the standard format used by NeuroMorpho.org (Ascoli, Donohue, and Halavi 2007), to date the largest collection of publicly accessible 3D neuronal reconstructions. While the SWC format is sufficient for compartment modeling and basic morphometry applications, it offers minimal or no support to detailed subcellular structures such as spines, somas, varicosities, and puncta, as well as support for other biologically relevant structures such as blood vessels or for integrated annotations constructs such as markers, text, regions, and contours. It also lacks the means to preserve information about any original source data.

Over the last twenty years, a shift from analyzing slides at the microscope to capturing images and image volumes for post-hoc analysis has steadily occurred. This change has expanded annotation potential for microscopy modalities (fMOST, microCT, EM, multiconfocal, lightsheet) and image data sizes. The advancement of microscope technologies with increasing resolution and imaging capabilities together with exquisitely specific labeling methods (pseudorabies transsynaptic labeling and transgenic techniques to name a couple), has enabled the visualization of smaller and subtler structures. This has driven the need for representing the relationship between neuronal morphology and smaller, subcellular structures such as spines, synapses, and varicosities or boutons (Jacobs et al. 2001; Le Bé, Silberberg, Wang, and Markram 2007; Arellano, Benavides-Piccione, DeFelipe, and Yuste 2007).

Starting in 2007, MBF created a new file format based on the Extensible Markup Language (XML). XML, a World Wide Web Consortium (W3C; 2008) standard, is well accepted and recognized as having a number of important attributes for data sharing: simplicity, extensibility, and self-description. The new format, often referred to as the Neurolucida XML format, is the subject of this article. It recreates the same data elements already present in the ASC and DAT files, while greatly improving data accessibility and extensibility. Additionally, XML’s hierarchical structure allows the contained reconstruction data to unambiguously denote the relationship of described elements in the context of their anatomical region at the tissue or organ level.

Very recently, a collaboration with the FAIR Data Informatics (FDI) lab through participation in the NIH Common Fund Initiative, Stimulating Peripheral Activity to Relieve Conditions (SPARC), has prompted MBF Bioscience to adapt the XML file format once again to embrace Findable, Accessible, Interoperable, and Reusable (FAIR) data standards (Wilkinson et al. 2016). The project highlights the importance of generating FAIR digital reconstruction and modeling data for anatomical and neuronal structures by incorporating the necessary metadata at the file level. MBF Bioscience has taken steps to ensure the image segmentation data produced by its products abides by the FAIR data standards for neuroscience, defined and promoted by the International Neuroinformatics Coordinating Facility (INCF; Wilkinson et al. 2016). As demonstrated by implemented data standards such as the Neurodata Without Borders neurophysiology data standard (Rübel et al. 2019), defined data format standards help to ensure generated resources are reusable and reproducible by the scientific community. We believe that embracing the transparency and systematic organization of the XML data file format will permit for further accommodation of the scientific community’s advocacy for open and accessible research.

## Purpose

The MBF Bioscience neuromorphological segmentation file structure has been driven for over 30 years by the ever-advancing science, technology, and input of neuroscientists throughout the world. The purpose of this paper is to document the relevant, systematic, and flexible nature of the file structure and demonstrates its effectiveness as a solution for the microscopic anatomy morphology data standard. Through these definitions, the neuromorphological file specification serves as an important format for the exchange of neuromorphological data for archival and exploratory research. We hope this will facilitate the development of other software and tools for downstream application and investigation of the increasing scope of data that is stored in this file format.

## File Structure Summary

A brief description of selected data elements of the neuromorphological file structure are detailed in this section. These elements were chosen to highlight their unique data structure’s direct impact on microanatomical models, analytics, and data reusability. The neuromorphological file structure is fully defined in the Neuromorphological File Specification (4.0) available at http://www.mbfbioscience.com/filespecification (Angstman et al. 2020). The file specification will continue to be updated as needed to define added and/or modified data elements in use with related MBF Bioscience software.

The neuromorphological file format is an <u>Extensible Markup Language (XML) 1.0 (Fifth Edition)</u> format and includes two organizational aspects, elements, and attributes. These aspects are defined in the XML file specification provided by W3C (The World Wide Web Consortium 2008). All neuromorphological data files include header elements, introducing the file via essential metadata, followed by morphological structure modeling elements.

## File-Level Metadata

In the header section, metadata attributes define the expected file structure, the software application, and version number of the neuromorphological file structure. By establishing the expected data structure for the document, the ability to interpret provisional file formats is preserved. Header elements are not used to model morphological structures and therefore differ from traced data elements.

Essential information regarding the software application includes the application name, application version, application Research Resource Identifier (RRID), and institution RRID. The institution RRID specifies the company, organization, or institution that produced the software which generated the neuromorphological file. Both application RRID and institution RRID are globally unique and persistent identifiers registered in the SciCrunch knowledge base. Reporting metadata regarding the software application used to generate the data file ensures the data generated is reproducible, reusable, and citable. This metadata will never get separated from the traced data because it is all saved at the file-level, unlike a sidecar metadata file.

### SciCrunch InterLex Terminology Linkage

Another header section of the neuromorphological file format stores critical subject and annotation metadata for each data file. The subject metadata is user-defined (left panel of Fig. 2a) and includes fields for subject species, identifier, sex, and age of the sample. The anatomical terminology list used for annotating neuromorphological or fiducial structures are selected using an API connection with the SciCrunch InterLex Terminology Portal (right panel of Fig. 2a). This information recorded in the <atlas> child element of the <sparcdata> section of the neuromorphological file format (Fig. 2b). The metadata accommodates organ, species, parcellation, and International Resource Identifier (IRI) to the atlas or ontology database if no atlas is available.

**Fig. 2.**
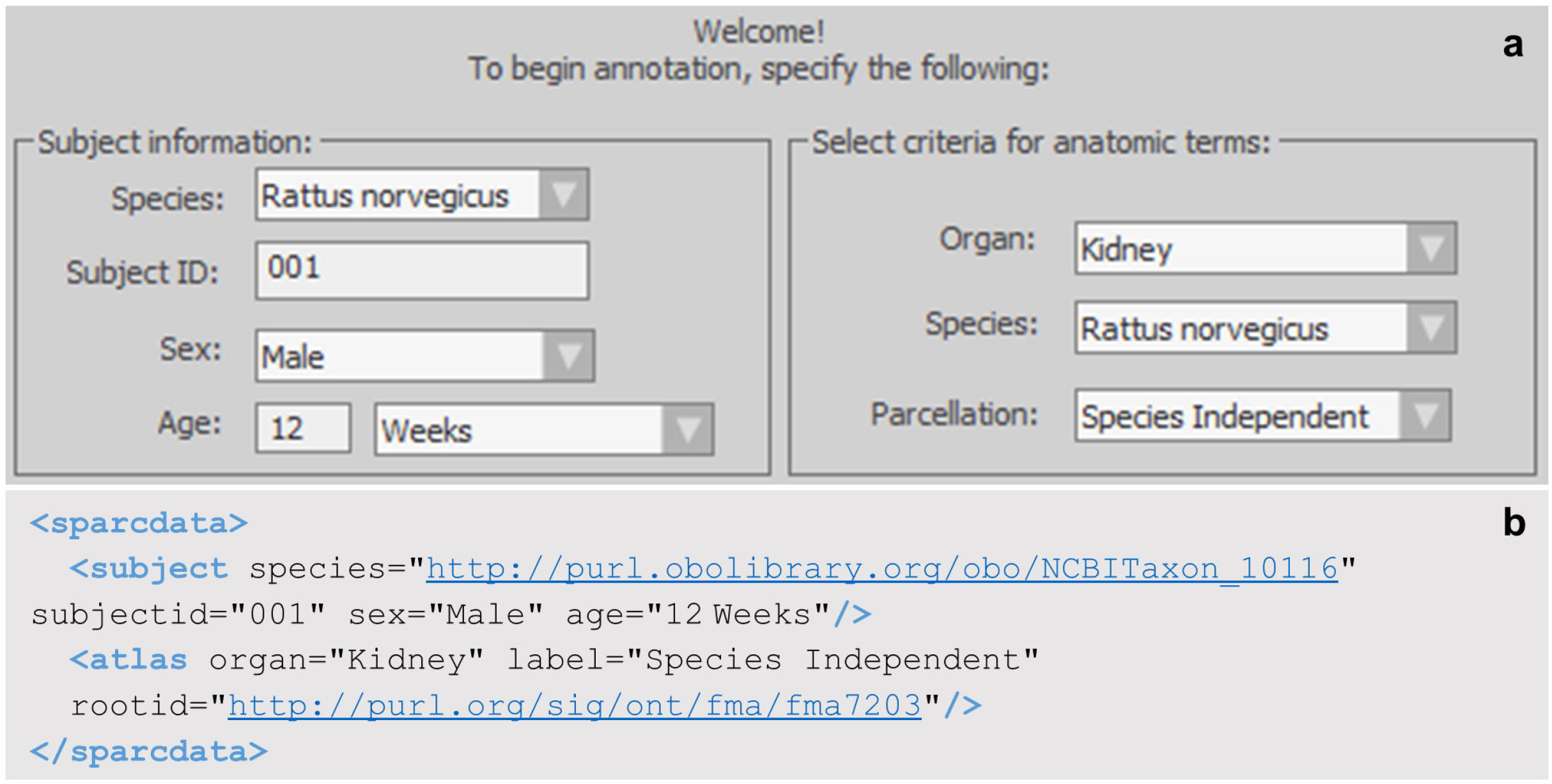
**(a)** The subject and annotation term list selection window within MBF software. The fields in the left section detail the subject information of the sample origin. The fields in the right section determine the anatomical term list provided to the user for annotation. The selected values indicate this sample originated from a 12-week-old male rat with the subject identifier 001. The anatomical term list selected for annotation was the rat kidney term list. The parcellation indicates Species Independent, meaning there currently is no parsed term list for the rat kidney. Instead, a generic term list of all kidney anatomies, independent of species, is provided to the user. **(b)** A neuromorphological data file related to a microscopy sample from a 12-week old male rat kidney delineated using anatomical terminology from the Foundational Model of Anatomy (FMA) ontology database. The species and <atlas> rootid store IRIs that are linked to the species and the term list origin selected in Fig. 2. The IRI includes the unique identifier for the species or parcellation.

The unique identifiers for the subject species, the anatomical term list, and delineated anatomical regions are recorded in the data file. This aids in repurposing and reusing any anatomically relevant data. A list of terms separated by organ, species, and atlas/parcellation scheme is provided to the user for annotation. Terms in the SciCrunch database are managed by SciCrunch (Grethe et.al. 2014). This infrastructure allows investigators to use a consistent and structured lexicon when referring to multi-species and multi-scale anatomies. With the API connection to the database, up to date term lists can be provided for annotation, ensuring the data file produced provides a robust understanding of how the traced data was derived.

SciCrunch services link recognized synonyms to the term identifier guaranteeing a database query of a term abbreviation or synonym will still return all applicable results by pulling any data tagged with the term. The clear file-level subject and annotation term list metadata within the neuromorphological data files can be queried and sorted by species, subject ID, sex, age, and organ of the sample origin.

### Coordinate space

The schematic in Fig. 3 demonstrates the coordinate space for all neuromorphological data files is three-dimensional. All coordinates and measurements are reported in micrometer units (μm). The origin point of the coordinate system is (0, 0, and 0).

**Fig. 3.**
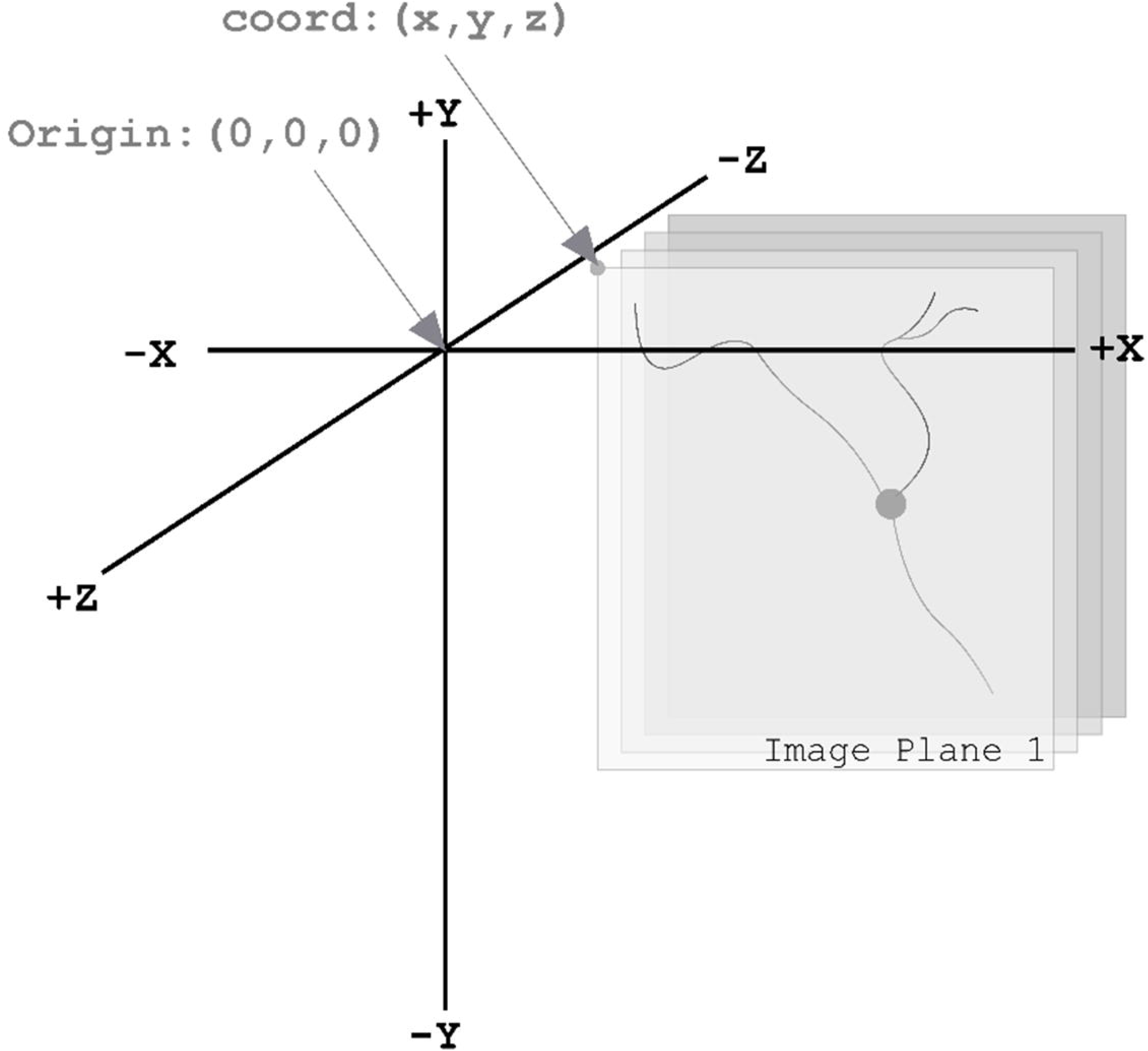
Demonstrates the 3D coordinate space with an origin point of (0, 0, 0). The gray planes represent a 3D image volume with an image location coordinate, coord: (x, y, and z). Note the direction of the Z-axis. The most positive image plane of the 3D volume is the first image plane. The following image planes are in the same X and Y location, but their Z location changes incrementally based on the z scaling. The units of this coordinate space are in micrometers (μm).

Using the image scaling defined as micrometers/pixel, the anatomical structures are readjusted into a real-world coordinate space enabling robust qualitative and quantitative analysis to be performed on the data elements. The example in Fig. 4b demonstrates the digital reconstruction of a 3D microscopy image (Fig. 4a). The morphometric measurements in Table 1 were obtain using the reconstruction in Fig. 4b and Neurolucida Explorer’s neuronal summary analysis (MBF Bioscience 2020).

**Fig. 4.**
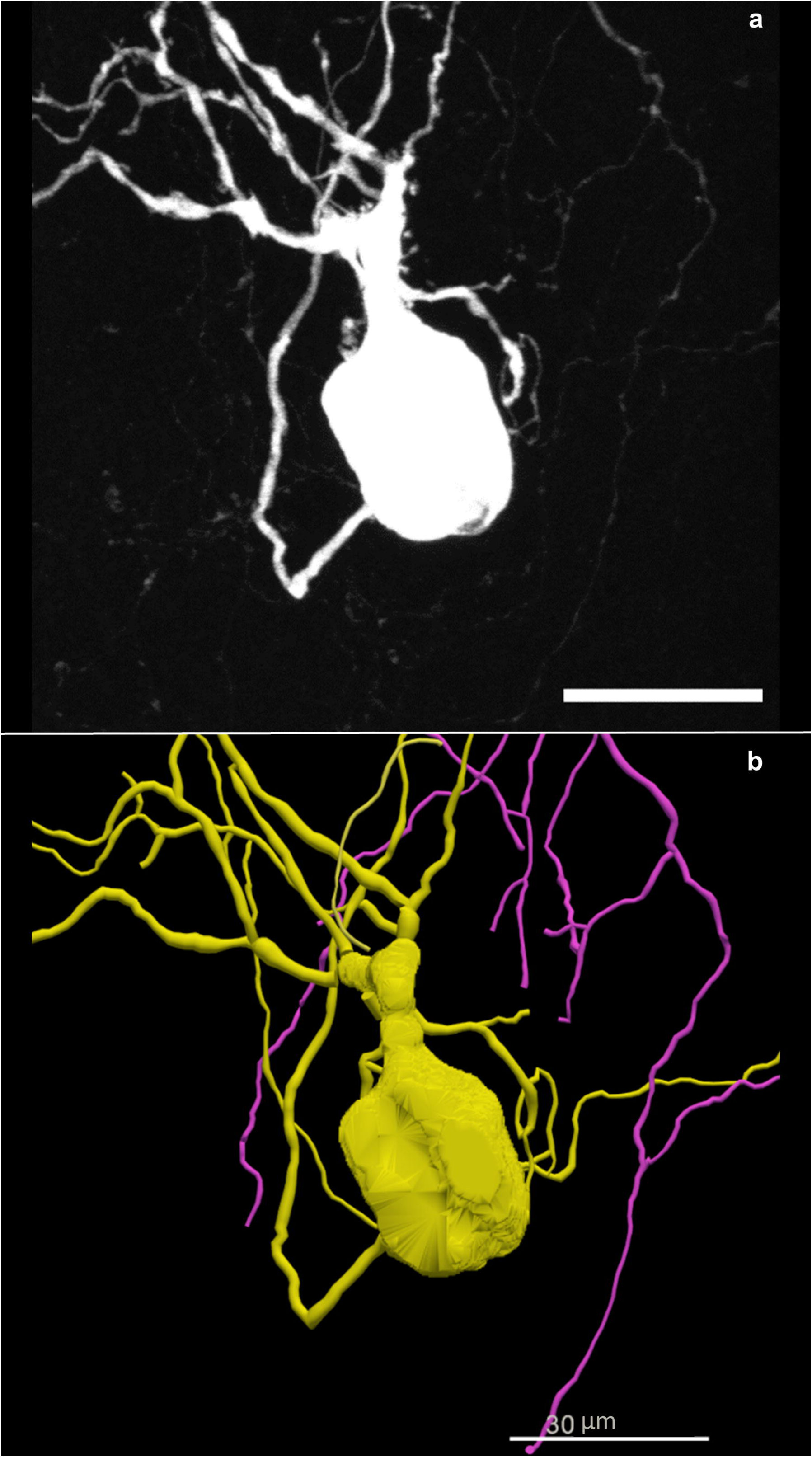
**(a)** A neuron from the mouse stellate ganglion backfilled with Neurobiotin. A 3D image was acquired on a Leica confocal microscope with a 40x objective lens. The scale bar equals 30 micrometers. The image scaling is: X= 0.2228 μm/pixel, Y= 0.2228 μm/pixel, Z= −0.5 μm. **(b)** A tracing of the neuron in Fig. 4a created using Neurolucida 360. The trace data elements include contours that make up the cell body and trees that reconstruct the neuron’s dendrites and axon. The scale bar equals 30 micrometers (Cho et al. 2020).

**Table 1.** Neuron Summary Analysis. The Neuron Summary Analysis produced from the tracing of the cell in Figure 3b. The length reported for cell body indicates the perimeter of all contours that make up the individual cell body (MBF Bioscience 2020). Length, mean length, surface, mean surface, volume, and mean volume are reported for contour and tree elements. The area and mean area analysis is only applicable for the 2D contours that construct the cell body. The value is reported as N/A (not applicable) for Axon and Dendrite trees. The complexity refers to the normalization and comparison of trees among fundamentally different neurons (Pillai 2012). This analysis does not account for cell bodies, therefore, N/A is the value reported.

| Name | Length<br>($\mu\text{m}$) | Mean<br>Length | Area<br>($\mu\text{m}^2$) | Mean<br>Area | Surface<br>($\mu\text{m}^2$) | Mean<br>Surface | Volume<br>($\mu\text{m}^3$) | Mean<br>Volume | Complexity |
| --- | --- | --- | --- | --- | --- | --- | --- | --- | --- |
| Cell<br>Body | 1440.628 | 72.031 | 6135.247 | 306.762 | 14407.086 | 14407.086 | 1992.254 | 1992.254 | N/A |
| Axon | 1013.302 | 1013.302 | N/A | N/A | 3180.255 | 3180.255 | 897.261 | 897.261 | 1013.302 |
| Dendrite | 1395.549 | 232.591 | N/A | N/A | 4659.897 | 776.649 | 1450.534 | 241.756 | 43262.016 |

**Table 2.**
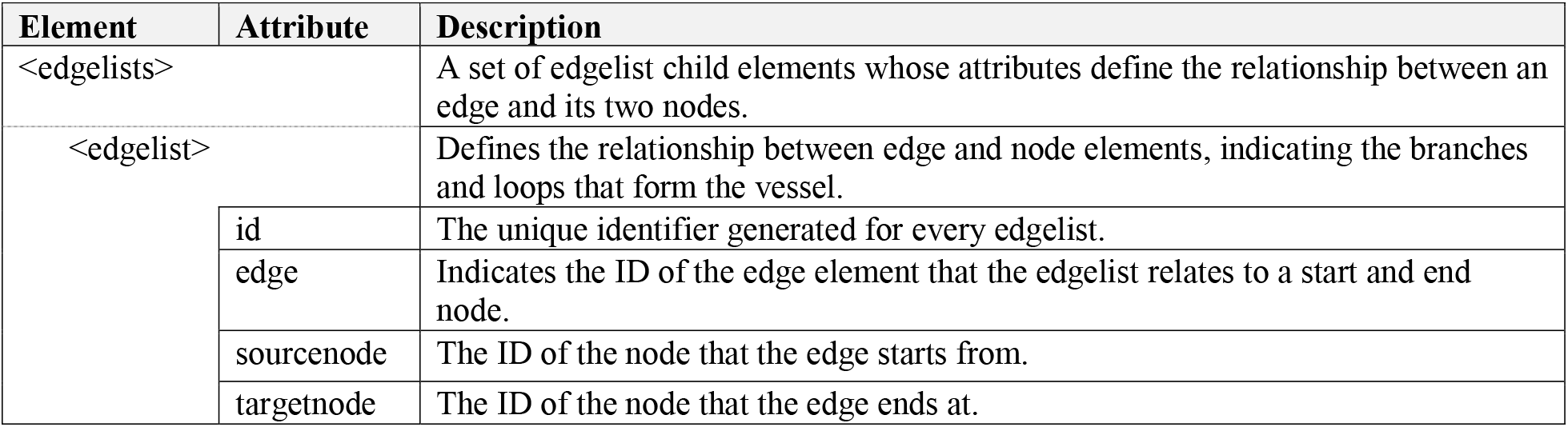
A description of child elements and attributes of the neuromorphological file’s edgelists element. A description for the edgelists child element of a vessel, its child elements, attributes, and values. An indentation before the element name and a dashed boarder is used to indicate a child element in the table.

As seen in the Table 1, the individual data elements (cell body, dendrite, and axon) can be analyzed as unique structures though they are all part of the same neuron. The importance of storing detailed coordinates for each individual element within the neuromorphological data file lies in the analysis flexibility. Not only can these structures be analyzed individually, but they can be analyzed in relation to one another. Each dendritic tree can be broken down to determine the length and volume of each tree. Using the same data, the volume of dendritic trees that fall within an anatomical volume such as an airway or a colonic layer can be determined, providing the neuronal density within that region.

### Microscopy Image Association

Raw image data is not saved within neuromorphological data files, rather they are linked with a file location, name, and other information about the image. Because the image data is not saved to file, the tracing data file is “light weight”, easy to store, transfer, and read. However, the file path and name of the image(s) are conserved, conveying the provenance of the data as it relates to the images in which it was derived.

The neuromorphological data file can be associated with any number of source images. The images can be either 2D (a single image plane) or 3D (multiple image planes from a single file or multiple files). Image data can be combined in several ways inside the neuromorphological data file. The simplest is the single 3D or 2D image file. The lineage of the derivative data is recorded in the file, regardless of complexity. This type of record keeping is rarely catalogued in other digital reconstruction data formats file formats.

Image scaling in X and Y (micrometers per pixel) and the Z scaling (micrometers between image planes) for the image(s) is reported alongside the related image name and file path. Additionally, the total number of image planes is included to further convey the associated image(s) structure. The X, Y, and Z coordinates of the upper, left-hand corner of an image is also reported. These values provide the location of the image within the data file coordinate space. To reuse multi-image neuromorphological files, it is necessary to know the image scaling, number of image planes, image order, and image location within the 3D coordinate space. These elements are stored alongside the corresponding image file path(s) and name(s). The file-level metadata indicating all source image(s) associated with the morphological data promotes the reusability of the data captured within the neuromorphological file format.

The ability to relate neuronal morphology reconstructions to multiple source images is a valuable aspect of the neuromorphological file format’s versatility. An application of this feature includes the generation of morphological and anatomical reconstructions on 2D and 3D image montages. In the case that multiple 2D image planes are combined to construct a 3D image volume, all image paths and file names are recorded in the neuromorphological file. The image order and Z location, key metadata for repurposing this data, is noted alongside the image element for these image types. The same is true for a singular image generated by merging source images that make up color channels to construct a multi-channel microscopy image. Lastly, registration of neuronal reconstruction from a high-resolution image to an annotated low-resolution image is possible due to this structure. This multi-resolution image segmentation process can provide anatomical context, especially in large organs or tissue samples that are difficult to image comprehensively at high magnification.

This case study depicted in Fig. 5 demonstrates how Cho et al. utilized multi-resolution image segmentation strategies to understand the structure-function relationships of cells within the stellate ganglion (Cho et al. 2020). After performing electrophysiological recordings, the cells were labeled with a fluorescent dye and processed for imaging. Using Neurolucida 360, Cho et al. reconstructed select neurons from images with a 40x objective lens (Fig. 5b). The entire stellate ganglion was also imaged with a 10x objective lens (Fig. 5c) and contoured using integrated FAIR anatomy terminology lists. The neuron reconstruction of the 40x, high-resolution image (Fig. 5b) was registered to the appropriate location in the 10x image (Fig. 5f demonstrates the x, y, and z location of the backfilled cell bodies can be discerned), providing anatomical context within the stellate ganglion. The registration of cellular reconstructions to the whole stellate ganglion allowed the researchers to bring the physiological and morphological data into context with the entire ganglion (Cho et al. 2020).

**Fig. 5.**
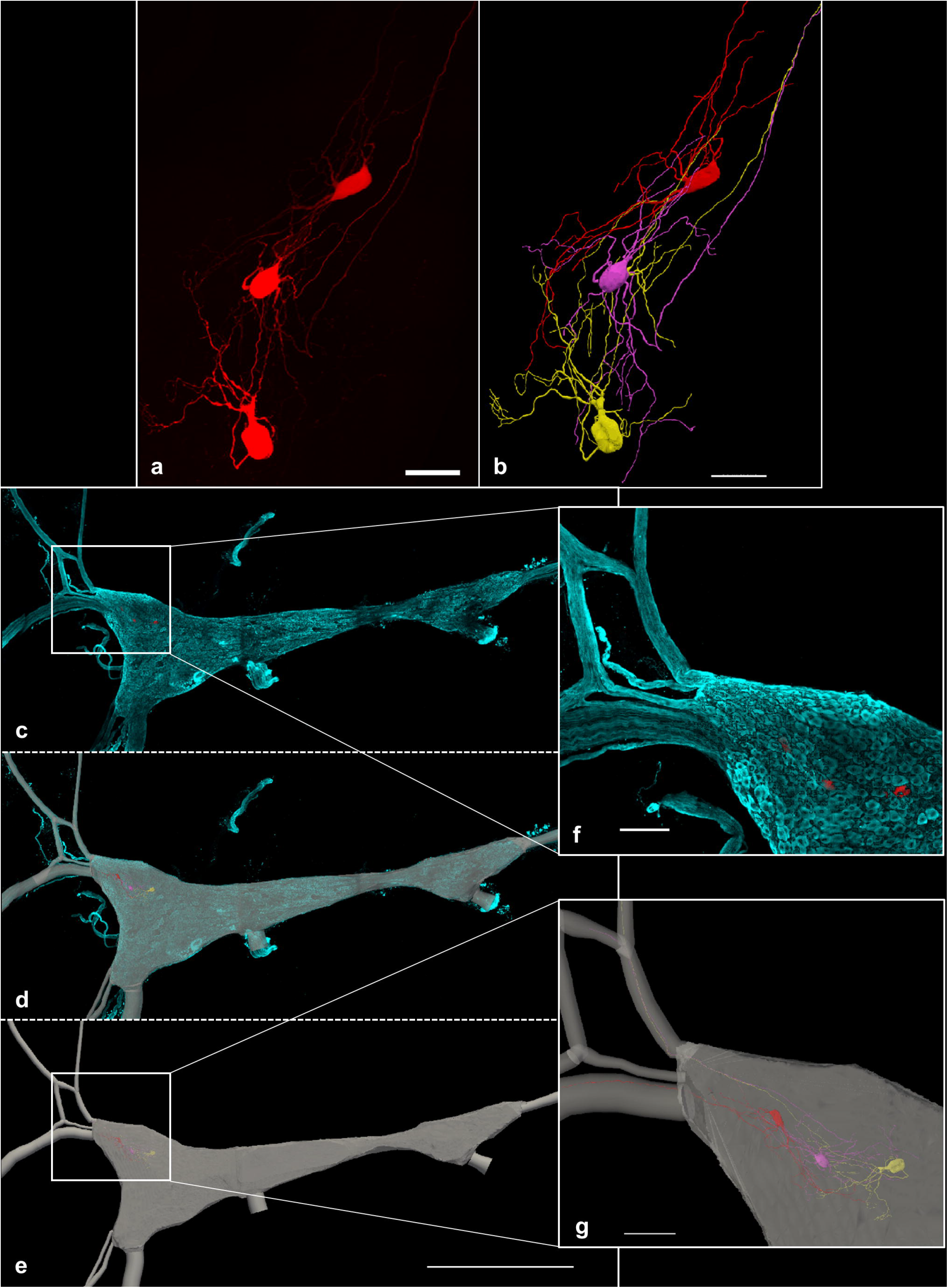
This figure illustrates the application of the multi-resolution image segmentation that is possible with the neuromorphological file format. **(a)** The 3D image, acquired on a Leica confocal microscope using a 40x objective lens, shows neurons from the stellate ganglion backfilled with Neurobiotin. The scale bar equals 50 micrometers. **(b)** A neuronal reconstruction obtained from the 40x, high-resolution image in Fig. 5a. Tree elements were used to represent the neuronal dendrites and axons of the cells. The cell bodies are represented using serial Z contours, shelled into a three-dimensional volume. **(c)** A 10x, low-resolution tile scan image was acquired using Leica confocal microscope. The images include the entire stellate ganglion labeled with tyrosine hydroxylase (TH) (cyan) and the neurons backfilled with Neurobiotin (red). This same group of backfilled neurons were imaged at 40x (Fig. 5a). **(d)** A 3D reconstruction of the 10x, low resolution whole stellate ganglion image (Fig. 5c) overlaid with that image. **(e)** 2D contours of the ganglia’s area were delineated at serial z image planes. They were shelled into a 3D volume to represent the stellate ganglion (gray). Tree elements were used to represent the path of the nerve fibers stemming from the ganglion. These structures were segmented using the 10x image in Fig. 5c. The 1000 micrometer scale bar shown in Fig. 5e is applicable for Fig. 5c, Fig. 5d, and Fig. 5e. **(f)** A zoomed in snapshot of the boxed location displayed over Figure. 4c. Scale bar equals 100 micrometers. **(g)** A zoomed in snapshot of the boxed location displayed over Figure. 5e. Axon innervation to the Inferior cardiac nerve (top) and Ventral ansa subclavia (bottom) can be mapped and visualized. Scale bar equals 100 micrometers (Cho et al. 2020).

Incorporation of file-level metadata including microscopy image association is an essential component of the data provenance of the morphology file. The neuromorphological file format takes the approach of maintaining the original source image as its original format, ensuring all source image metadata is conserved (e.g. objective magnification, channel IDs, modality). By incorporating access to both the source image and its derived digital reconstruction files, data repositories can enable user to re-use, repurpose, or reproduce this data by providing all necessary components. By only saving the file location(s) and name(s), factors like image dimensions, number of image planes, number of channels, and bit depth do not affect the reconstruction data file size. Storing this data can increase the size of the reconstruction data file by over 99%. File size can bog down downloading, 3D visualization, and web rendering speeds. For those only interested in repurposing the reconstruction data, this is an exceptional benefit as it notably decreases load times. These factors illustrate the value of the neuromorphological file format’s microscopy image association scheme.

## Morphological Structure Modeling

The traced data elements include all data models of neuromorphological structures and additional annotations. A data file will not necessarily contain all types of traced data elements and typically will include more than one of a single traced element in a data file. For example, one file could have cell body contours and neuronal trees where another could incorporate cell body contours, neuronal trees and marker elements.

A variety of trace data elements have been added to the neuromorphological file, their format augmented for performance, morphometric modeling accuracy, and analytical potential based on feedback from top neuroscientists in the field. Below, the structure of relevant neuromorphological data elements are detailed providing the background for discussing the FAIR aspects of the data and the structure’s relevance to representing and analyzing the neuromorphology.

### Trees

In the neuromorphological file structure, the tree element is used to represent non-looping branching structures within microscopy data such as axons, dendrites, and airways. Trees consist of an origin, branches, nodes, and endings. The one neuronal dendrite within the 3D confocal image show in Fig. 6a was reconstructed (Fig. 6b, Fig. 6c). The reconstructed dendrite is represented in the schematic in Fig. 6d to demonstrate the neuromorphological file’s tree data structure. The starting point of a tree element is referred to as the origin (O) and the points that follow make up the root branch of the tree. All trees must have at least an origin and root branch, but typically have branching points called nodes. Nodes are where a segment of the tree splits into multiple branch child elements. The branch elements are made up of an ordered list of points that connect nodes to nodes, and nodes to endings. Endings are the last point of a branch or tree where the segment terminates.

**Fig. 6.**
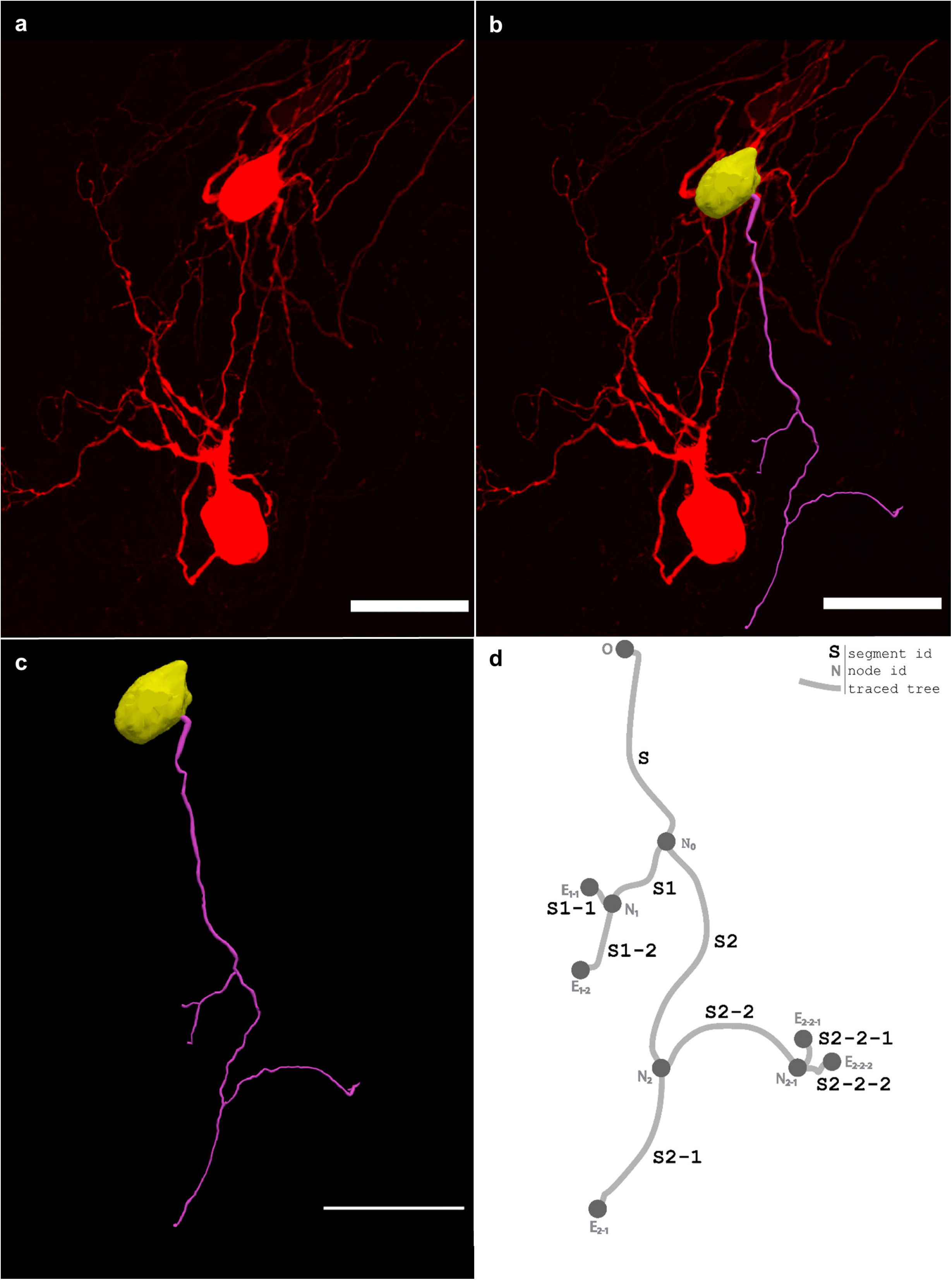
**(a)** The 3D image, acquired on a Leica confocal microscope with a 40x objective lens, shows neurons from the stellate ganglion backfilled with Neurobiotin. **(b)** A 3D reconstruction of one dendrite and one cell body of the backfilled neuron from the stellate ganglion (Fig. 6a) overlaid with that image. **(c)** The same reconstruction shown in Fig. 6b with no image data. A model of one cell body (yellow) and one neuronal tree (pink) was produced using Neurolucida 360 (Cho et al. 2020). **(d)** A diagram demonstrating the structure of a tree with each segment shown as a line and labeled with the segment name (ex. S2-2-2). The origin (O), nodes (N) and endings (E) of the tree are marked with a circle. The root segment (S) begins with the origin (O) point and terminates with the node (N_0_). The child segments of N_0_, S1 and S2, terminate with nodes N_1_ and N_2_. The child segments of N_1_, S1-2 and S1-1 terminate with endings E_1-1_ and E_1-2_. N_2_ has two child segments, S2-1 and S2-2. Segment S2-1 has no bifurcations, so it terminates with ending E_2-1_. Segment S2-2 bifurcates at node N_2-1_. Lastly, the branches S2-2-1 and S2-2-2 terminate with endings E_2-2-1_ and E_2-2-2_. All scale bars are equal to 50 micrometers.

The neuromorphological file format stores each point location (x, y, z position, and diameter) for a tree and its branches. The tree element is formatted as described above, to clearly relate relevant anatomical features such as the nodes indicating origin, ending, and bifurcation points. High level and detailed analysis can be performed on the computer readable morphometric reconstruction data of trees and branches. This includes quantifying the number of branches or terminals within an anatomical region or determining the proximity of two branches such as a neuronal fiber and a blood vessel. By classifying the neuronal fiber types, for example axons, dendrites, and apical dendrites, the total number of each can be determined and the details of the trees can be compared. Tree details including length, surface area, volume, total terminations, branch angles, etc. Even the complexity and extension of neurons can be determined (Table 1, Column 9). Analyzing realistic, meaningful, and quantifiable neuron reconstructions can help researchers to conclude the structure-function relationship that defines a specific subset of neurons.

Reporting each point of the tree enables child element to be associated at unique locations along the branch. This is valuable for representing neuronal morphologies such as spines, synapses, and varicosities. By representing the relationship of structures like the tree and spine in the data file, joint analysis can be performed determining the tree’s spine density, spine class densities, or average spine diameter. Even the distance from an individual spine’s base to the tree origin can be calculated. The spine-tree association is further detailed in the section below.

### Dendritic Spines

Spines are small protrusions off of dendritic branches. The neuromorphological data format embeds the spine element in the associated branch at the tree points that the spine occurs.

The Backbone property of the spine describes the points that construct the spine volume (Fig. 7a). The number of total points is listed first. Following this, the X, Y, and Z coordinates along with the diameter of each coordinate are listed in order of proximity to the branch. The diameter thickness of each point determines the three-dimensional shape of the modeled spine. The first point is the branch insertion point along the branch centerline. The next two point model the base and neck of the spine. The fourth and fifth points model the head and tip location of the spine.

**Fig. 7.**
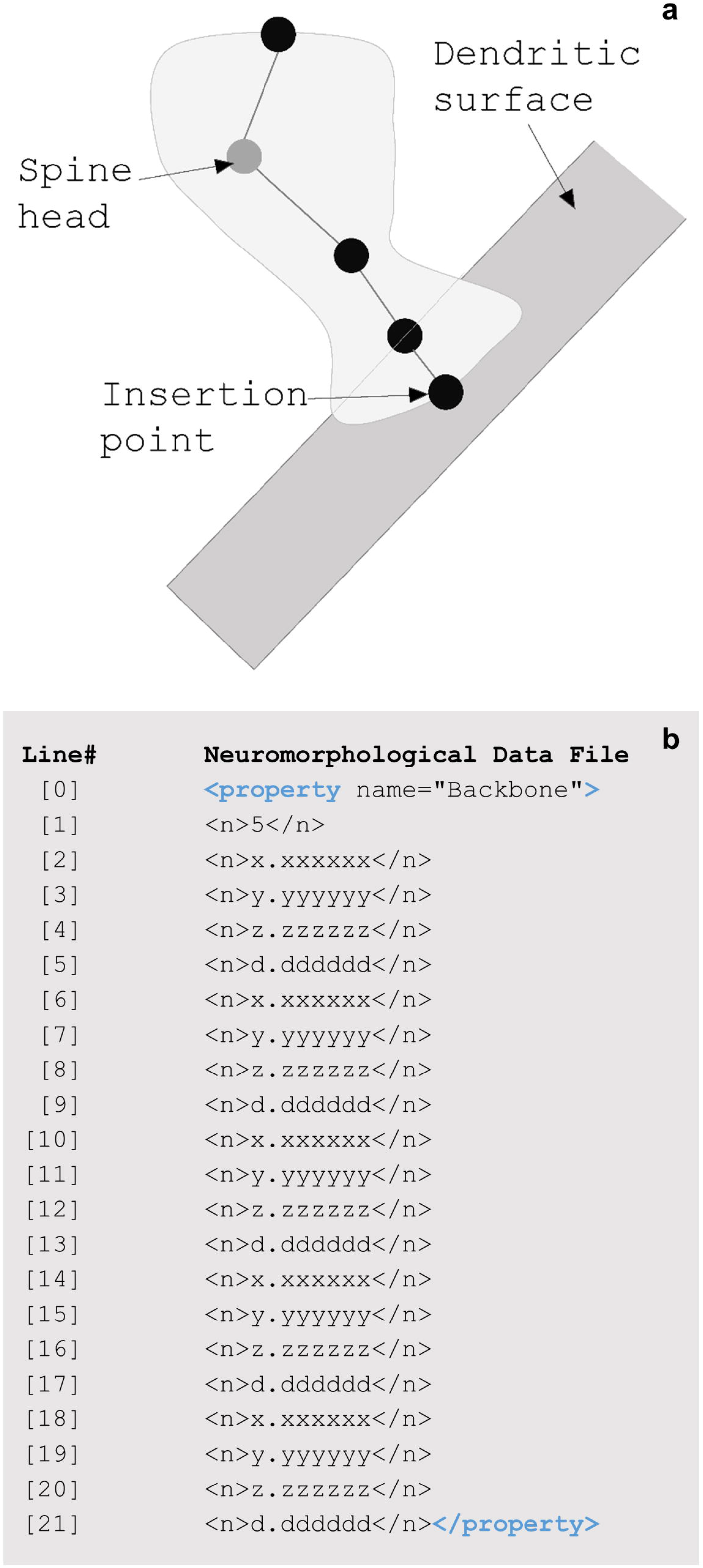
**(a)** A diagram of a dendritic spine along a neuronal tree. The five points of the spine are represented with circles. The coordinates of these points are reported in the <property name=”Backbone”> number string including an x, y, and z location along with a thickness, d. The spine head is marked with a gray circle. **(b)** The Backbone of a spine includes a string of numbers. The line numbers and return spaces present in Fig. 7b were added for clarity and do not exist in the data file structure. Line [1]’s value reports the total number of points that make up the spine. The values from line [2] through [21] make up each of the four spine coordinates (x, y, z, and d). The first point (x=line [2], y=line [3], z=line [4], and d=line [5]) listed is the insertion point where the spine is located along the tree.

By modeling the key dynamic connection component of brain cells, dendritic spines, we can analyze how these dendritic spines grow, disappear, and change shape over time. Researchers are able to extract accurate 3D measurements of these dendritic spines to validate novel techniques testifying to the data’s reliability. By analyzing 1,500 dendritic spine reconstructions, Gao et al. were able to confirm that physically increasing tissue size through expansion microscopy did not cause damage to the structural components. This helped to demonstrate the utility of the novel expansion microscopy (ExM) technique and validate its ability to obtain comprehensive morphometrics of delicate dendritic spines in combination with lattice light sheet microscopy (LLSM) and digital reconstructions. Due to the data’s format, the range of spine head diameters, neck diameters, backbone lengths, and neck backbone lengths could be extracted from the morphometric models. The team found the spine metrics collected from the expansion lattice light sheet microscopy (ExLLSM) samples proved consistent with a relevant electron microscopy (EM) study (Gao et al. 2019). As mentioned above, the relationship between spines and dendritic trees can be analyzed to provide a contextual understanding of the nanostructure’s arrangement on a unique grouping of neuronal fibers. Individual spine metrics written to the data file include but are not limited to volume, classification (Rodriguez, Ehlenberger, Dickstein, Hof, and Wearne 2008), and backbone lengths. Even subunits of the spine can be compared such as spine head position, head length, and head diameter vs. neck length, and neck diameter. The level of detail available for analysis is due to the file format’s detail of the element used for modeling spines.

### Anastomoses

Structures like vasculature, nerves, or fascicles, can be represented with a different branching structure than neuronal trees. In the neuromorphological file format, vessel elements are made up of points called nodes that connect branches called edges (Fig. 8a). An edge is a collection of connected points. The connection relationships are described with the edgelists element (Fig. 8b).

**Fig. 8.**
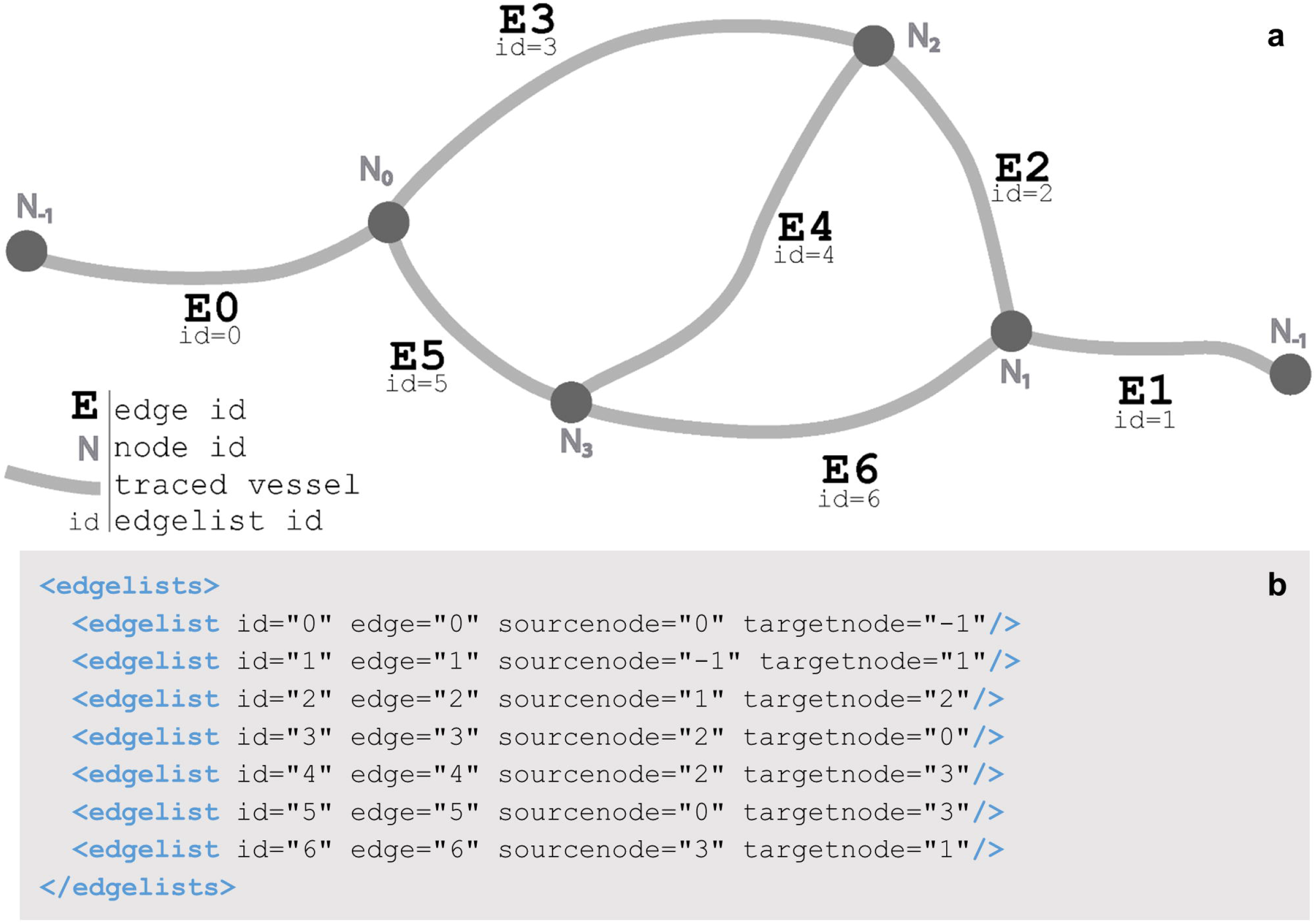
**(a)** A diagram of the edgelists element of Fig. 8b. Each edgelist and edge id correspond to one of the vessel branches. These are labeled appropriately. The edgelist sourcenode and targetnode inform the start and endpoint of the vessel branch or edge. For example, edge=”4” (E4) begins at node 2 (N_2_) and ends at node 3 (N_3_). This connection is indicated in edgelist id=4 (see Fig. 8b). This connection of the vessel back onto itself creates a loop structure. **(b)** The data structure for the edgelists child element of a vessel. Each edgelist id attribute corresponds to the edgelist ids in Fig. 8a, informing how the vessel edge elements connect to the node elements. The edge attributes correspond to the edge ids in Fig. 8a. The sourcenode and targetnode values refer to a node id in Fig. 8a.

The vessel elements can have edges that loop, whereas tree elements can only branch. The looping capability of the vessel elements is modeled with what is known as a graph structure. The graph structure has a wide range of applications and can be used to model anything from hyperlinks in webpages to transportation of goods (Siek, Lee, & Lumsdaine 2001). What makes it so powerful for modeling anatomical morphologies is its ability to represent biological structures such as anastomoses in vascular or nerve networks.

Due to the data structure that stores each unique vessel point and its diameter, detailed analysis of length, surface area, volume, average thickness, and even tortuosity can be obtained through the vascular models. A recent study performed vascular reconstructions were generated of micro-CT rat brains on control and blast-exposed rats to examine the effects of traumatic brain injury on vascular networks (Gama Sosa et al. 2019). Though the micro-CT images showed a clear decrease in vasculature structures, the reconstruction of both control and blast-induced rat brains provided meaningful quantitative results to back this observation. The results showed that the blast-exposed rat brains decreased in length by about 50%, in surface area by about 50%, and in volume by about 60% (Gama Sosa et al. 2019).

The structure also permits anastomose analysis including loop length, surface area, volume, and average diameter, which can aid in the classification of looping structures. The data can guide researchers in predicting the function of each loop class based on its morphometric traits.

Like other data elements of the neuromorphological file, vessels can be analyzed at a high level, reporting the network summary, loop count, or the branching within an anatomical structure. By coupling two data types, vessels and contours, we are able to further our understanding of vasculature abnormalities that may occur. For example, reconstructing anatomical regions alongside vasculature networks can allow for the exploration network loop density within specific regions of control and treatment samples. More applications of utilizing contours for delineating anatomical regions in two or three dimensions is further detailed in the following section.

### Anatomical Structure Delineation

A contour element is a named list of sequentially connected points. Contours are often used to delineate anatomical regions within image data. The TraceAssociation property of the contour stores an IRI that is linked to the anatomy term used for a contour and includes the unique and persistent identifier for that term. If this child element is present in the contour, then the contour name and TraceAssociation are both saved to the data file via the API connection with the SciCrunch InterLex Terminology Portal discussed in the above section. The contour name is selected from the anatomical term list recorded in the <sparcdata> <atlas> element. The contour element in Fig. 9b has the name “Glomerulus” which was selected from the species independent kidney terminology list. The corresponding metadata associating this term to its term list is exemplified in the <atlas> element of Fig. 9b.

**Fig. 9.**
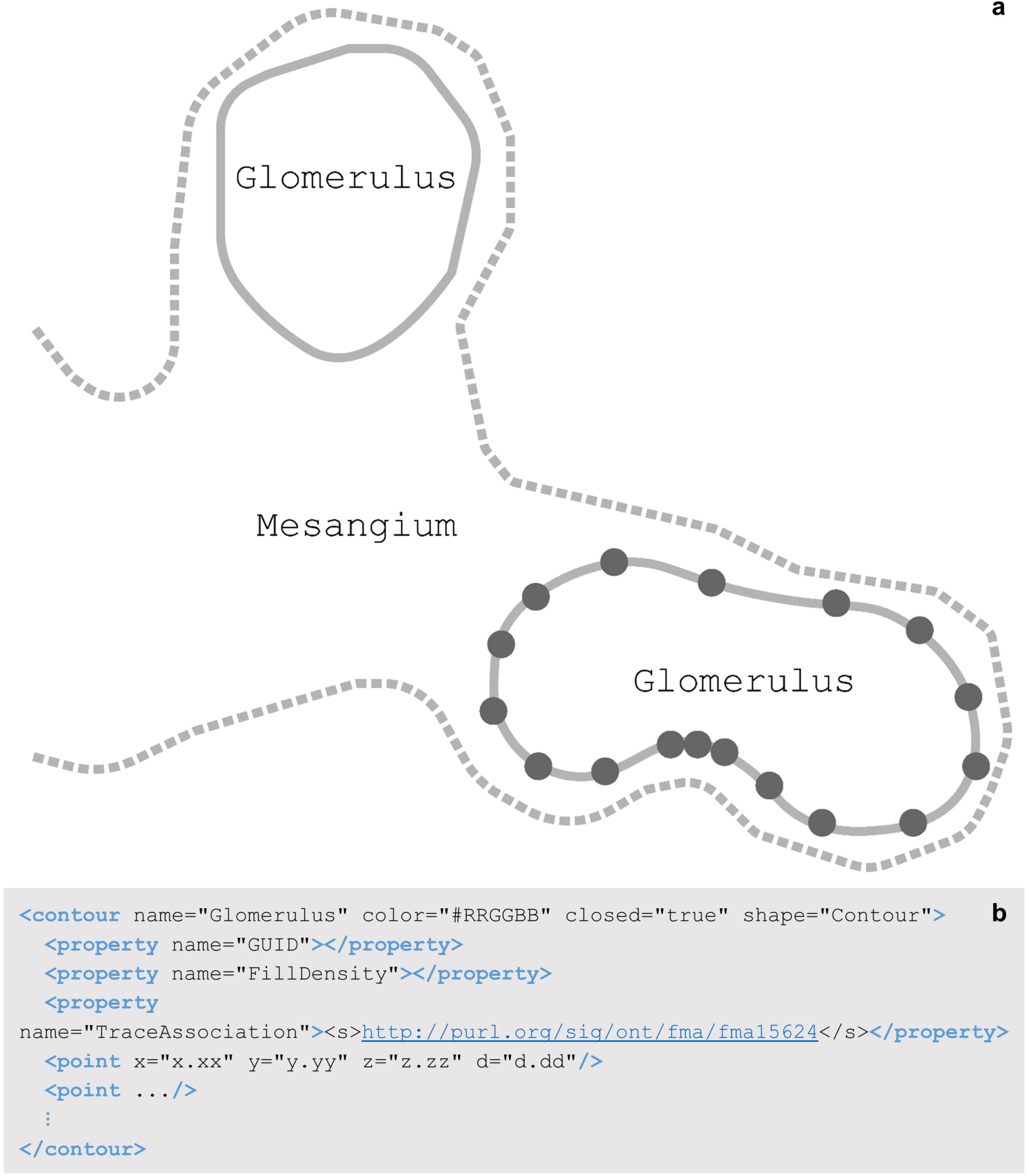
**(a)** A schematic of a contoured regions of a renal corpuscle. The marked point locations (represented as circles on one glomerulus contour) are connected with a line (solid, dashed) to generate an area that represents an anatomical region in two-dimensions. The dashed line represents the mesangium region where the solid lines represent glomeruli. The glomeruli contours are closed where the mesangium is an open contour indicating the structure continues and the contour represents the layer of the mesangium that falls within the renal corpuscle. **(b)** A Glomerulus contour element, child elements, attributes, and values as they appear in the segmentation data file. This contour is a closed contour indicating the first and last point elements are connected. In this Fig. 9b, the <property> child elements exclude all values for concision with the exception of the TraceAssociation property. The value of the TraceAssociation property is the IRI to the Glomerulus term in the FMA kidney ontology term lists. The point elements have been abbreviated in this Fig. 9b. A contour usually contains a list of many point elements, connected in the order they are listed in the contour.

2D or 3D cell bodies are represented using the contour element or groups of contour elements. All cell body contours have names containing “soma”. 3D cell bodies are traced using multiple contours of the same name at different Z positions outlining the entire Z space of the cell body region.

Contours can be used to delineate 2D structures or they can be grouped to represent a 3D surface or volume. For example, 3D cell bodies are traced using multiple contours of the same name at different z positions, outlining the entire Z space of the cell body region. These structures can provide anatomical context or they can be used in data analysis. The density of a reconstructed neuronal network within an anatomical region can be analyzed based on the volume of the region’s contours.

In a recent study by Achanta et al., researchers utilized the neuromorphological file format to delineate regions of the rat heart and mark the location of neurons of the intrinsic cardiac nervous system (ICN). The contour element was used extensively to map the cardiac structures in two dimensions (Fig. 10a) – such as the endocardium of left and right atria, auricles, and ventricles – within the histological sample to create a 3D reconstruction of the heart (Fig. 10b; Achanta et al. 2020). The annotation methodology was used again by Leung et al. to compare ICN neuron location and distribution across multiple male and female rat hearts (Leung et al. 2020).

**Fig. 10.**
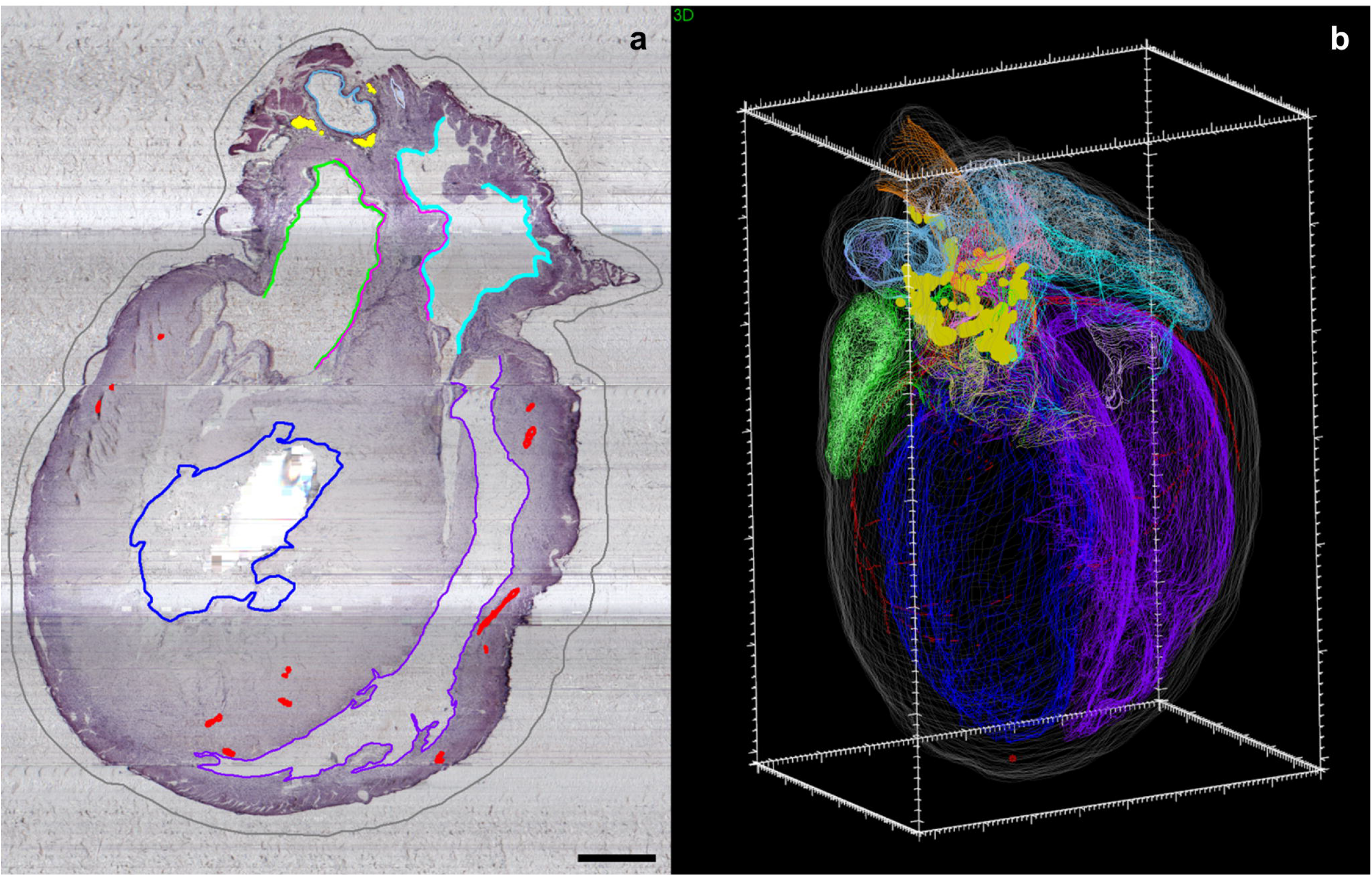
The anatomy of the male Fischer rat heart, including the intrinsic cardiac nervous system neurons (yellow), were mapped using MBF Bioscience’s TissueMapper application **(a)** to create a comprehensive 3D reconstruction **(b)**. **(a)** The scale bar is equal to 1000 micrometers. **(b)** The 3D scale bar’s minor ticks are equal to 100 micrometers on the x-axis, 200 micrometers on the y-axis, and 100 micrometers on the z-axis.

Contour-style segmentation can be performed in other software applications and widgets, such as ImageJ’s ROI Manager Tool, but the output file format (e.g., ROI file) often lacks data provenance, structural organization, and is unreadable to humans outside of the ImageJ application. The contour element within the neuromorphological file format addresses these concerns with clear data lineage, concise and human-readable detail regarding the point makeup of each contour as it relates to the structure within the microscopy image(s), and is packaged in a single file for further ease of reuse and interoperability.

### Point Markers

The marker element is used to represent single point locations in the data file. This is useful for mapping and/or counting cell types (Zaborszky et al. 2015), synapses (Le Bé, Silberberg, Wang, and Markram 2007), and other punctuate objects. Additionally, they can be used to mark fiducial points or anatomically significant single coordinate locations, as seen in Fig. 11, to assist the mathematical registration of segmentation data to a generic 3D scaffold model.

**Fig. 11.**
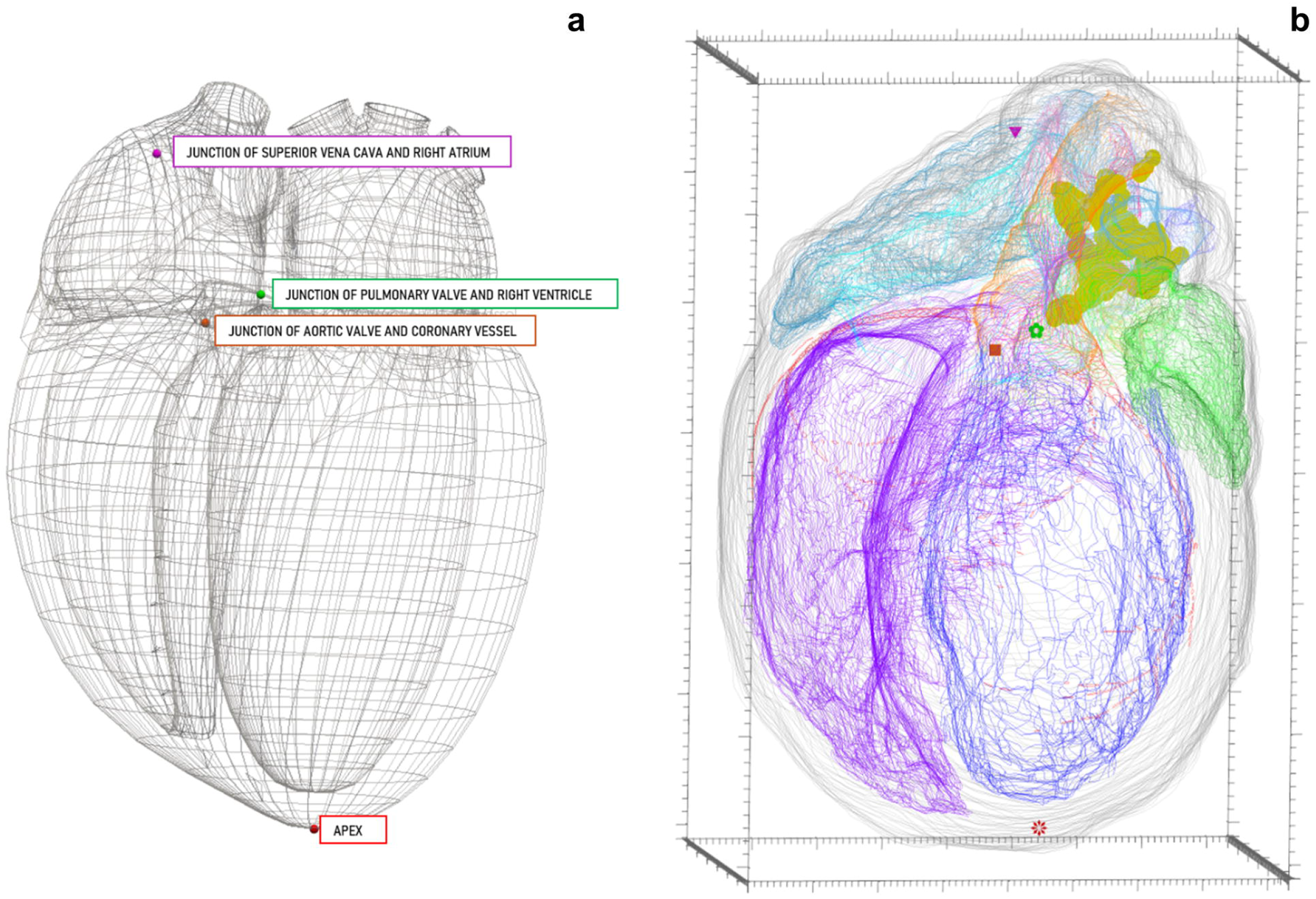
A generic heart scaffold **(a)** beside segmentation **(b)** in a multi-viewport 3D environment to identify and mark concordant fiducial points essential for registration with the common coordinate scaffold. The triangle marker in Fig. 11b represents the discrete location of the junction of the superior vena cava and the right atrium. This location is also marked in Fig. 11a in the matching marker color and alongside the associated marker name. The following marker pairs follow the same format as described above for the triangle marker: flower marker (Fig. 11b) = junction of the pulmonary valve and the right ventricle (Fig. 10a), square marker (Fig. 11b) = Junction of aortic value and coronary vessel (Fig. 11a), star marker (Fig. 11b) = Apex (Fig. 11a). **(b)** The 3D scale bar’s minor ticks are equal to 100 micrometers on the x-axis, 200 micrometers on the y-axis, and 100 micrometers on the z-axis.

Since the marker element can be used to represent a variety of biological and non-biological structures, there are a myriad of ways to classify the markers within the data file, allowing others to properly decipher a data file with little inquisition. These attributes include name, color, and shape, all of which can be applied to multiple or individual marker elements.

One application of marking fiducials for registration to the 3D organ scaffolds engineered by the Auckland Bioengineering Institute (ABI; Leung et al. 2020). These scaffolds allow researchers to compare segmentation data from multiple subjects by matching fiducial points in each segmentation file to those same fiducial points within a generic organ scaffold. Mapping segmented data to a common space can help correct for sample deformation due to experimental imaging protocols. By fitting the samples to the generic organ scaffold via fiduciary point matching, objects of interest can be compared to each other with true anatomical context.

### Sets

Set properties can be found in any trace data element. It is utilized for naming and grouping data of one or many trace data elements. These elements can either be the same type (e.g., all tree elements) or different types (e.g., spine, tree, and contour elements). By placing traced elements into a set, researchers can embed their knowledge and expertise on the sample into the traced data file, bridging gaps between those reusing and repurposing the data. Applications for the set property include: defining relationships to anatomies, annotating anatomies consisting of more than one traced data element, associating traced elements and supplemental data, and grouping objects to select and edit all at once. For example, neurons can be grouped together with a set name “intraganglionic laminar endings” to represent a muscular afferent within the colon, a key neuronal structure for sensing muscle stretching. This annotation is carried with the segmentation data and can be incorporated in downstream applications and investigations. Elements can also be associated with multiple set properties. This is one of the set property’s most useful abilities because annotations about an overarching structure or relationships to other data modalities can be made while still including anatomical context for those structure. Fig. 12c demonstrates one use case for associating multiple sets with one data element. Note the multiple set properties indicating this axon is associated with the electrophysiology readings number 18105039-091 and the axon innervation. Those looking to repurpose this data file can identify which of the dataset’s electrophysiology files pair with this axon. The anatomical context of the axon within the greater structure of the stellate ganglion is apparent though the set property denoting the path of the fiber.

**Fig. 12.**
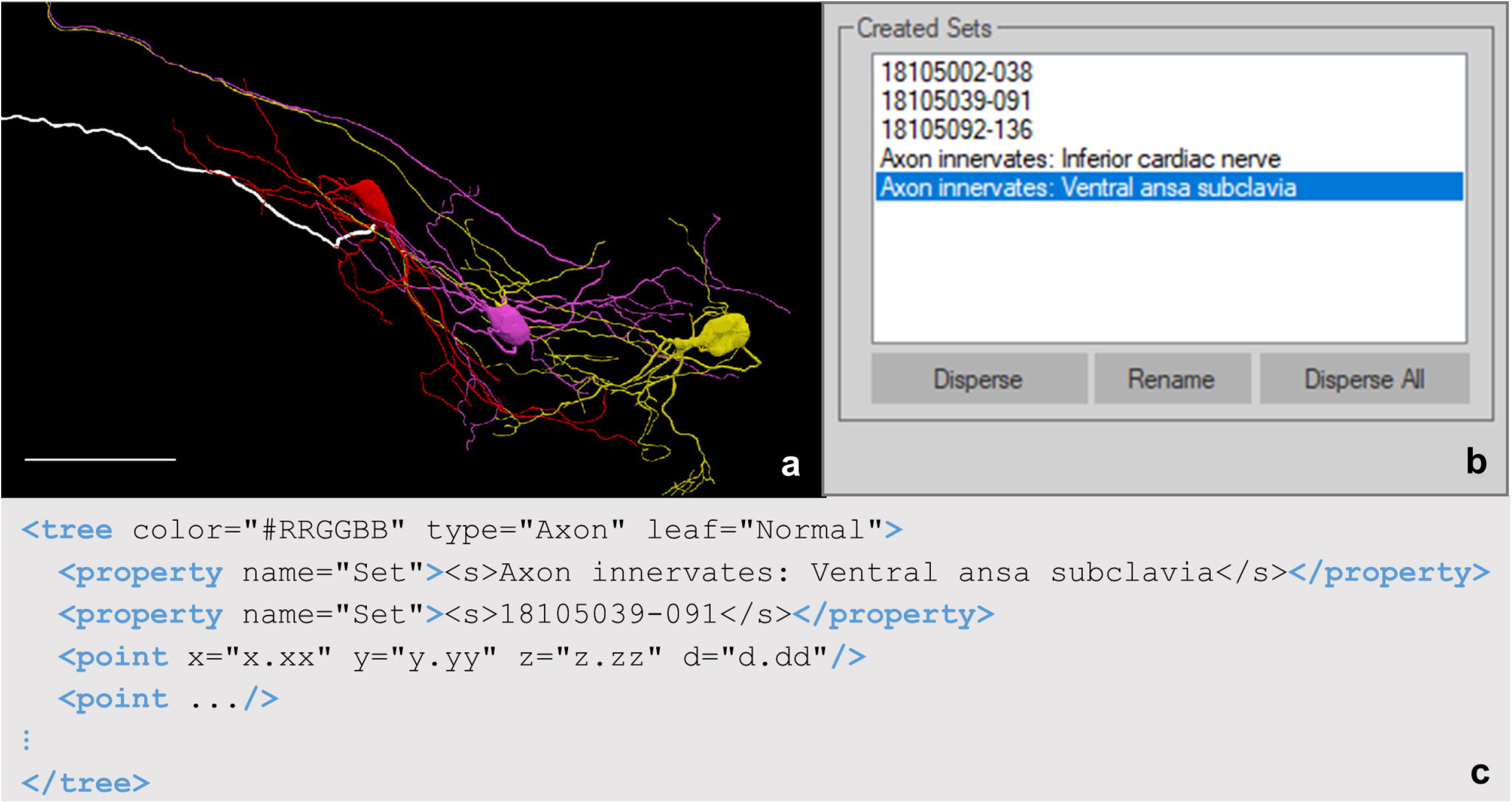
**(a)** Neuron reconstructions generated with Neurolucida 360 based on the image described in Fig. 5a. Highlighted in white is the axon for cell 18105039-091. The cell ID corresponds to the electrophysiology readings taken for each cell backfilled with Neurobiotin. The scale bar is equals to 100 micrometers (Cho et al. 2020). **(b)** The names of all created set in the tracing shown in Fig. 12a. The highlighted set, Axon innervates: Ventral ansa subclavia, describes which nerve of the stellate ganglion cell 18105039-091 innervates. **(c)** The tree element of the axon for cell 18105039-091. The point elements in this tree have been abbreviated using an ellipsis to draw focus to the structure of the set properties of the axon.

**Fig. 13.**
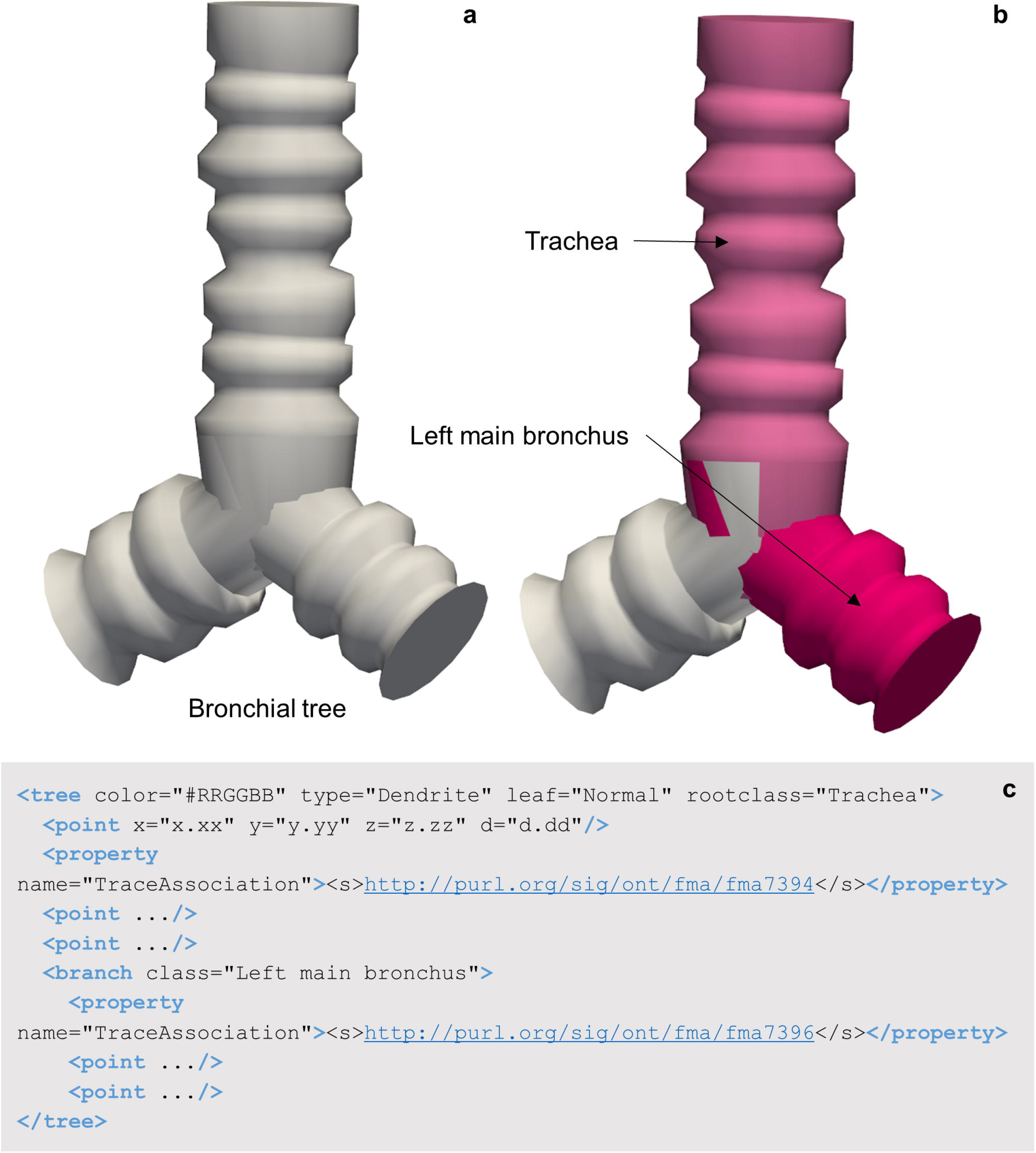
**(a)** A 3D reconstruction of a bronchial tree utilizing the tree elements of the neuromorphological file format. **(b)** The Trachea and left main bronchus segments of a lung airway named using the SciCrunch terminology link through MBF Bioscience software. **(c)** The data representation of the bronchial tree. Segment 1 (S1) of the bronchial tree is classified as the Trachea. The branch point indicates S2 of the bronchial tree is classified as the Left main bronchus. Unique identifiers for each term are indicated in the segment’s Trace Association property.

The data collected by Cho et al. demonstrates the importance of the set property in communicating additional relationships through the neuromorphological file format. Each backfilled cell from the stellate ganglion was reconstructed (Fig. 12a; Cho et al. 2020) and placed into a set named with the electrophysiology identifier (Fig. 12b). The axon innervation was labeled for each cell indicating the nerve fiber that the axon passes through as it exits the stellate ganglion. Encapsulating this otherwise autonomous anatomical context known only by original researchers promotes easy and constructive reusability of this file format’s modeled morphologies.

## Discussion

The neuromorphological segmentation file structure has evolved for over 30 years. It is used in software applications such as Neurolucida and Neurolucida 360, which has become prevalent systems for neuron reconstruction, with more than 6,500 citations. The file format is widely utilized in projects including the Human Brain Project, the Blue Brain Project, and the NIH SPARC program. Currently, the Blue Brain Portal hosts over 1000 Neurolucida neuron reconstruction data files, and the Human Brain Project’s EBRAINS Knowledge Graph has accumulated around 100 Neurolucida files to date (Retrieved August 6^th^, 2020). Additionally, other software e.g., NEURON, a simulation environment for neurons and networks can utilize Neurolucida data files for computational modelling applications. Publication of the file format specification will increase the utility and sustainability of neuromorphological data across research fields and continue to support data-driven science, while aiding in data management.

The structural elements of the neuromorphological file format were influenced by input and feedback from leading neuroscientists to propel scientific discovery using quantifiable digital reconstructions of biological structures. The structural elements were also designed to generate Findable, Accessible, Interoperable, and Reusable (FAIR) data and metadata. The scientific community continues to move toward an open and collaborative climate. FAIR data standards are being developed and adopted by funding institutions, further influencing the shift. Another factor driving FAIR data is the growing importance of bioinformatics and database queries due to the surplus of scientific data of all kinds. Due to the growing emphasis on producing FAIR data, we plan to submit the neuromorphological file format as a standard for digitally reconstructed microscopic anatomies referred to as the neuromorphological file standard. The elements of the neuromorphological file were designed to be Findable by humans and computers to allow the file type to be queried by online databases via globally unique, persistent, and up-to-date identifiers. These identifiers are searchable via the SciCrunch InterLex Terminology Portal. The file-level information within the neuromorphological files can be leveraged by common fund research projects or initiatives (i.e., HuBMAP) to enhance searching functions within online collectives of experimental data. The human readable and published file format is Accessible to anyone looking to repurpose this data type (Angstman et al. 2020). Encoded in the well-recognized XML format, the metadata and tracing data’s Interoperability is one of the neuromorphological file format’s vital attributes (World Wide Web Consortium 2008). The detailed specification of the file format is open to the public. Its elements have been decided to enhance interoperability with limited human interaction. Because the format is XML and human readable, it can be easily view and parsed with a variety of software, e.g., MATLAB and Python, extending its outreach to drive scientific research forwards. Tools such as NEURON, the neuronal simulation modeling software, already support Neurolucida reconstruction data. By publishing the neuromorphological file specification we hope to expand the utility of this data type and encourage re-use and repurposing of the reconstructions. The metadata conserved in the header elements of the neuromorphological file provides a robust understanding of how the traced data was derived such as the generating software, sample origin, and image scaling. The association of the morphometric modeling data with this metadata provides file-level information on the data’s origin. Because this metadata is stored to the same location as the morphology models, the two are never separated ensuring the Reusability of these data types. These characteristics demonstrate that the neuromorphological file format meets the FAIR data principles developed and endorsed by the INCF (Wilkinson et al. 2016). However, there are many additional and unique benefits to each element of the neuromorphological file. These include the thousands of morphometric analyses available for each distinct element or for elements in combination. These relevant advantages of the neuromorphological file format validate the file format as a rich digital reconstruction format for microscopic neuronal anatomies.

A broadly accepted, neuronal tracing format, such as the neuromorphological file format, that is open and FAIR can catalyze scientific research (Meijering 2010; Halavi, Hamilton, Parekh, and Ascoli 2012; Parekh and Ascoli 2013).

## Governance

The neuromorphological file format specification is open to the community and can be accessed at http://www.mbfbioscience.com/filespecification. The file specification will continue to be updated as needed to define added or modified data elements. To facilitate the integration of community contributions, we plan to generate a standard mechanism for enhancements such as documentation corrections, data storage, and feature requests. The bases of the community contributions will be facilitated through the publicly available web forum generated to provide support for tool developers implementing the file format. We refer interested participants to https://forums.mbfbioscience.com/.

## Code availability

Due to intellectual property right restrictions, we cannot provide source code or its documentation for the commercially available Neurolucida, Neurolucida 360, Vesselucida 360, and Tissue Mapper software. Free trials are available at http://www.mbfbioscience.com.

## Future Directions

Currently, annotators utilizing MBF Bioscience’s software link to the SciCrunch InterLex Terminology select curated vocabulary terms and segment that anatomical region of interest utilizing the contour element. Building upon this functionality, we plan to establish a FAIR system for linking anatomical terms and unique identifiers to tree, vessel, and marker data modeling elements. This addition will allow researchers to readily conform to FAIR data principles for every annotated data element of the neuromorphological data format.

Tree structures such as neuronal branches or bronchial trees (Fig 13a) can be modeled and classified via SciCrunch curated ontologies. Fig. 13b demonstrates the classification of two branches of the airway with the appropriate anatomical names found in the SciCrunch term list. Note the data file (Fig. 13c) includes both anatomy terms (Trachea and Left main bronchus) and two separate term identifiers, one for each segment of the tree. Other anatomical structures that follow a non-looping branch pattern can use a similar modeling structure. Our goal is to incorporate this approach for linking vessels and marker elements with SciCrunch InterLex Terminology.

Following the publication of the neuromorphological file format, we propose to document the appropriate structure for linking microscopic modeling elements to the SciCrunch InterLex Terminology Portal. This effort will specify how to pull lists regularly to surface up-to-date anatomical ontologies. The user interface that specifies a select term lists will also be demonstrated as a guideline for other tool builders to implement this resource.

Lastly, we recognize the importance of community contributions to further enhance the neuromorphological file format. A plan is in place to generate an open community strategy that provides a structure submission mechanism for additions and modifications. Contact through email and forums will help provide the necessary checks and balances for this open and FAIR file format.

## Acknowledgments

We would like to dedicate this manuscript to the memory of Dr. Edmund Glaser in appreciation for devoting his career of more than four decades to the field of neuroscience. In 1963, he co-invented and patented the Image-Combining Computer Microscope and pioneered the method of quantifying 3D neuronal morphometry. This technology applied computer techniques to neuroanatomical research, permitting scientists to precisely quantify the brain’s three-dimensional structure. In 1987, Dr. Glaser co-founded the company MicroBrightField (now MBF Bioscience) with his son, Jacob (Jack) Glaser. Over the ensuing decades, reconstructing neuronal structures was reduced from hours to minutes and measurement precision was able to achieve sub-micron accuracy. Large assemblies of neuronal networks can now be examined in quantitative detail in three dimensions. Dr. Edmund Glaser’s accomplishments in the development of computer microscopy and his contributions to neuromorphological reconstruction were unparalleled and essential to the formulation of this file format.

We would also like to recognize Maryann Martone and the SPARC project for accelerating the publication of the neuromorphological file format.

## Author Information

### Affiliations

MBF Bioscience, Williston, VT, USA

### Corresponding author

Jacob R. Glaser

## Additional information

### Availability of data and material

The Neuromorphological File Specification (4.0) can be found at www.mbfbioscience.com/filespecification.

